# STAT3 inhibits Myocardin induced cardiac hypertrophy

**DOI:** 10.1101/2020.11.21.392902

**Authors:** Xing-Hua Liao, Yuan Xiang, Jia-Peng Li, Hui Li, You Huang, Chao Shen, Hui-Min Zhang, Tong-Cun Zhang

**Author notes:** Corresponding author: Prof. Tong-Cun Zhang, Institute of Biology and Medicine, College of Life Sciences and Health, Wuhan University of Science and Technology, Hubei, 430081, P.R.China., and Prof. Xing-Hua Liao, Institute of Biology and Medicine, College of Life Sciences and Health, Wuhan University of Science and Technology, Hubei, 430081, P.R.China. These authors contributed equally to this work (Xing-Hua Liao, Yuan Xiang).

## Abstract

**Background:** In order to explore the molecular mechanism of cardiomyocyte-dependent myocardial gene expression and cardiomyocyte differentiation in cardiac hypertrophy, and to provide new insights for cardiac hypertrophy.

**Methods:** Cardiac myocytes were isolated from day 1-3 Sprague-Dawley rat pups. Real time quantitative PCR, western blot and immunocytochemistry Assay were used to detect the expression and localization of related genes. CO-IP was used to detect direct protein interactions between Myocardin and STAT3. Luciferase reporter assay and chromatin immunoprecipitation were used to detect the binding of Myocardin to the promoter of a downstream target gene. Microinjection of zebrafish embryos was used to examine the effects of STAT3 and Myocardin interactions on cardiac development in vivo

**Results:** The N-terminus of STAT3 directly binds to the basic domain of myocardin and inhibits the transcriptional activity of Myocardin-mediated cardiac-specific genes ANF and α-actinin, thereby inhibiting their expression, and further inhibit myocardin-mediated cardiac hypertrophy in vivo.

**Conclusions:** In summary, our report states that signal transduction and transcriptional activation factor 3 (STAT3) are inhibitors of the major cardiac hypertrophic transcription factor Myocardiin, which is required for cardiomyocyte differentiation. The STAT3-cardiacin interaction identified nuclear hormone receptor-mediated and cardiac-specific gene-regulated convergence sites and suggested a possible mechanism for cardioprotective effects.

## Background

Myocardin play a key role in cardiovascular development and are specifically expressed in the heart and smooth muscle cells. (1) In addition, Myocardiin is a co-activator of serum response factor (SRF). SRF can be combined with the CArG box (CC [A/T] _6_GG), which is a DNA consensus sequence located in the control region of many growth factor regulated genes and muscle-specific genes.(2) Consistent with the role of Myocardiin as a transducer of hypertrophy signals, the forced expression of Myocardin in cardiomyocytes is sufficient to replace the hypertrophy signal and induce cardiac hypertrophy and fetal cardiac genetic programs. (3) In a word, Myocardin is a key component of molecular switches that regulate SRF’s ability to mediate cell proliferation and muscle cell differentiation. (4–6)

Signal transducer and activator of transcription (STAT) 3 is involved in cell survival, proliferation, and immune response.(7) STAT3 has been reported as a key mediator of cardiac remodeling in response to cytokines, especially glycoproteins (gp) 130 family, including cytokines of interleukin 6 (IL-6) and leukemia inhibitory factor (LIF).(8–10)

IL-6 binding results in receptor dimerization, which induces tyrosine phosphorylation and STAT3 activation through Janus-activated kinases. Activated STAT3 is then dimerized and transported to the nucleus to activate or inhibit downstream target gene expression. STAT3, as a key mediator of myocardial cell survival, seems to play an essential role in inducing myocardial hypertrophy. So far, there were not some researches about the effect of STAT3 in Myocardin-induced cardiomyocyte hypertrophy. This study reveals a relationship between Myocardin and STAT3 which are crucial transcription factors in cardiomyocyte hypertrophy development.

## Methods

### Cell culture

Cardiac myocytes were isolated from day 1-3 Sprague-Dawley rat pups as described previously.(11–13) COS7 cells were bought from American Type Culture Collection (ATCC) and cultured in Dulbecco’s modified Eagle’s medium (DMEM) (Invitrogen) supplemented 10% fetal bovine serum (FBS) in the incubator with a humidified atmosphere (5% CO2, 37°C).

### Plasmid construction and transfection

To overexpress Myocardin and STAT3, nucleotide fragments encoding the fusion Flag tag protein and full-length of Myocardin and encoding the fusion Myc tag protein and full-length of STAT3 were cloned to mammalian expression vector pCDNA3.1 (Invitrogen) (Flag-pCDNA3.1-Myocardin, Myc-pCDNA3.1-STAT3). Chemically synthesized small interfering RNA (Si-STAT3) and scrambled negative control (NC) Si-NC (Ribobio) to knockdown STAT3 expression. ANF-1000-luciferase reporter, ANF-337-luciferase reporter, ANF-236-luciferase reporter and ANF-121-luciferase reporter plasmid, ANF-1000-M-far-CArG, ANF-1000-M-near-CArG, ANF-1000-M-GAS, ANF-1000-M-far-CArG and GAS, ANF-1000-M-GAS and near-CArG and ANF-1000-M-far-CArG and near-CArG-luciferase reporter, were all constructed into pGL3-basic vector (Promega). All plasmid constructs were transfected into COS-7 cells or cardiomyocytes with FuGENE^®^ HD, and the experimental procedures were strictly in accordance with the manufacturer’s instructions.

### Immunocytochemistry Assay

Immunofluorescence assays were performed as described previously. (14) After added the primary antibodies (goat anti-rabbit ANF (Abcam), goat anti-mouse α-actinin (Sigma), and then added appropriate secondary antibodies (FITC-goat anti-rabbit IgG, FITC-goat anti-mouse IgG Santa Cruz), respectively. DAPI (4’,6’-diamidino-2-phenylindole) was used to stain nuclei, the samples were evaluated under inverted fluorescence microscope (Olympus).

### Reverse-transcription polymerase chain reaction (RT-PCR) and real time quantitative polymerase chain reaction (qPCR)

Methods for performing RT-PCR and qPCR analysis have been described previously.(14) Briefly, total RNA was isolated from cells using Trizol reagent (Invitrogen), the samples were reverse-transcribed by using M-MLV reverse transcriptase (Promega) according to the manufacturer’s instructions. The PCR primer sequences are as follows: Glyceraldehyde-3-phosphate dehydrogenase (GAPDH): F-GGAGCGAGATCCCTCCAAAAT, R-GGCTGTTGTCATACTTCTCATGG; ANF: F-CAACGCAGACCTGATGGATTT, R-AGCCCCCGCTTCTTCATTC; α-actinin: F-ATTGGTATGGAGTCTGCCG, R-TCCTGAGTGTAAGGTAGCCG; Myocardin: F-AGAACTCAGGGGCACACGAAG, R-CCACCTTGTCAGAAGATTGTAAACC. GAPDH was used as an internal control.

### Western Blot (WB)

The operation process of Western Blot has been described previously.(14) Cells were lysed in lysis buffer to extract total protein. The resulting protein was then analyzed on a 12% sodium lauryl sulfate polyacrylamide gel (SDS-PAGE). The proteins were transferred to a PVDF membrane and blocked for 60 min at room temperature in 5% skim milk powder (wt/vol) in TBST (TBS+0.1% Tween-20, vol/vol). After mouse polyclonal antibodies against Myocardin (Abcam), p-ERK (Abcam), rabbit polyclonal antibodies against ANF (Abcam), mouse monoclonal antibodies against α-actinin (Sigma) and appropriate secondary antibodies (HRP-goat anti-mouse IgG, HRP-goat anti-rabbit IgG, HRP rabbit-anti-goat IgG, Santa Cruz) were incubated. The specific proteins were visualized by odyssey detection. GAPDH expression was used as an internal control.

### Luciferase assay

The dual luciferase assay system (Promega) was used to measure luciferase activity and was standardized for transfection efficiency. Normalize results by dividing Firefly luciferase activity by Renilla luciferase activity of the same sample.

### Coimmunoprecipitation assay (CO-IP)

A plasmid-based expression vector encoding Flag-Myocardin and Myc-STAT3 was co-transfected into COS7 cells. Lysates were collected 48 h after transfection and Flag-Myocardin was precipitated using Flag antibodies. The resulting mixture was washed, subjected to sodium dodecy1 sulfate-polyacrylamide gel electrophoresis (PAGE), transferred to a polyvinylidene fluoride membrane, first developed by Myc antibody to visualize Myc-STAT3, then peeled off and again with Flag antibody Probe to reveal Flag-Myocardin. Myc-STAT3 was then precipitated using Myc antibody. The resulting mixture was washed, subjected to PAGE, transferred to a polyvinylidene fluoride membrane, first developed by Flag antibody to reveal Flag-Myocardin, then peeled off and re-examined with Myc antibody to reveal Myc-STAT3.

### Chromatin Immunoprecipitation (ChIP) assay

ChIP analysis was performed using an enzymatic chromatin IP (magnetic beads) kit (CST) in Cardiac myocytes transfected with Flag-Myocardin for 24 h. Proteins bound to DNA were crosslinked using formaldehyde at a final concentration of 1% for 20 min at room temperature. Protein-DNA complexes were immunoprecipitated using anti-Flag antibody (Santa Cruz) for Flag-Myocardin. The signal of Myocardin/ANF promoter complexes was measured by qRT-PCR. The primers used for amplification of the human ANF promoter were (ChIP-ANF-promoter-CArG1 forward: GCTGCTCAAGGCAAAGG) and (ChIP-ANF-promoter-CArG1 reverse: AGAGCTGGAACCCTCCC); (ChIP-ANF-promoter-CArG2 forward: ACCCACGAGGCCAATGAATC) and (ChIP-ANF-promoter-CArG2 reverse: GCTCCCTCTGCTTGCATCTCA). For the analysis of ChIP-qRT-PCR experimental results, calculate the enrichment relative to the input chromatin according to the delta Ct method with the percentages been calculated using the formula 2^-ΔCt^, where ΔCt is Ct (ChIP-template)-Ct (Input). A standard curve from one of the dilution inputs was included in the run to keep the reaction efficiency between 95% and 105%. qPCR was performed separately for each ChIP assay on control areas that should not bind myocardiin. The ChIP assay was considered specific if in the control region the enrichment was not above the enrichment of the non-specific immunoprecipitated sample made with normal rabbit immunoglobulin G.

### Zebrafîsh Strains and Microinjection

Dispose in accordance with local animal welfare regulations and maintain in accordance with standard procedures (www.ZFIN.org). The experiments were evaluated by the Animal Experiment Committee (DierExperimenten Commissie-DEC) and evaluated by the Animal Association Committee of Wuhan University of Science and Technology. The experimental procedure was in accordance with Directive 2010/63 / EU of the European Parliament on the protection of animals. The zebrafish strains used in this study were raised according to standard procedures. In order to knock down endogenous expression of Myocardin and STST3 in zebrafish, morpholinos (MO): Myocardin-MO, STST3-MO and control-MO targeting ATG(15) were synthesized from Gene Tools (Philomath, OR). Wild-type zebrafish embryo was injected at one-two cell stage with either Myocardin-MO, STST3-MO or control-MO. Zebrafish phenotype was analyzed under stereo microscope as previously described.(16)

### Whole-Mount Confocal Microscopy and Morphometric Analyses

In order to observe the shape and development of the heart, Whole-Mount Confocal Microscopy and Morphometric Analyses were used, as described previously.(16) Briefly, in order to examine the embryo heart, zebrafish embryos were anesthetized with 0.003% tricaine, place the ventral side of the embryo in the observation chamber towards the microwell, facing 45° to the right.. Confocal images were captured by using a Zeiss LSM 510 Confocal Microscope System. The confocal pinholes were adjusted to 1 Airy unit to achieve an optimal z resolution of 0.75 N.A. and obtain continuous slices of 0.9 μm thickness. Images were acquired at 0.45 μm intervals. 3D projections were constructed by using the LSM Browser software (Zeiss), and analysis was performed by Image J software that reads .l sm files (http://rsb.info.nih.gov/ij).

### Statistical Analysis

Data were expressed as mean±SE and accompanied by the number of experiments performed independently, and analyzed by t-test. Differences at *P*<0.05 were considered Significant differences in statistics.

## Results

### STAT3 antagonizes the activation of myocardial genes by Myocardin

Myocardin is reported as a key component of molecular switches that regulate SRF-mediated cell proliferation and myocyte differentiation. (1) STAT3 is a critical mediator for survival of cardiomyocytes and appears to be essential in the induction of cardiac myocyte hypertrophy. To explore whether STAT3 synergizes with Myocardin to promote cardiac hypertrophy, cardiomyocytes were transfected with Myocardin, STAT3 and Myocardin/STAT3 plasmids, and the mRNA and protein levels of cardiac genes were quantified by qPCR and WB. Our results indicated that overexpression of Myocardin significantly activated the expression of cardiac genes (ANF and α-actinin), while co-transfected Myocardin and STAT3 reduced cardiac gene expression induced by Myocardin (Fig. 1A-C). We also detected the ANF and α-actinin gene expression and localization by immunofluorescence, the result showed coexpession Myocardin and STAT3 can weak the Myocardin-induced cardiomyocyte hypertrophy (Fig. 1D, E). We also knocked down the expression of endogenous STAT3 using si-STAT3. Our results show that knocked down endogenous STAT3 can relieve the inhibition of STAT3 on the expression of Myocardin activation of ANF and α-actinin (Fig. 2A, B).

**Fig. 1.**
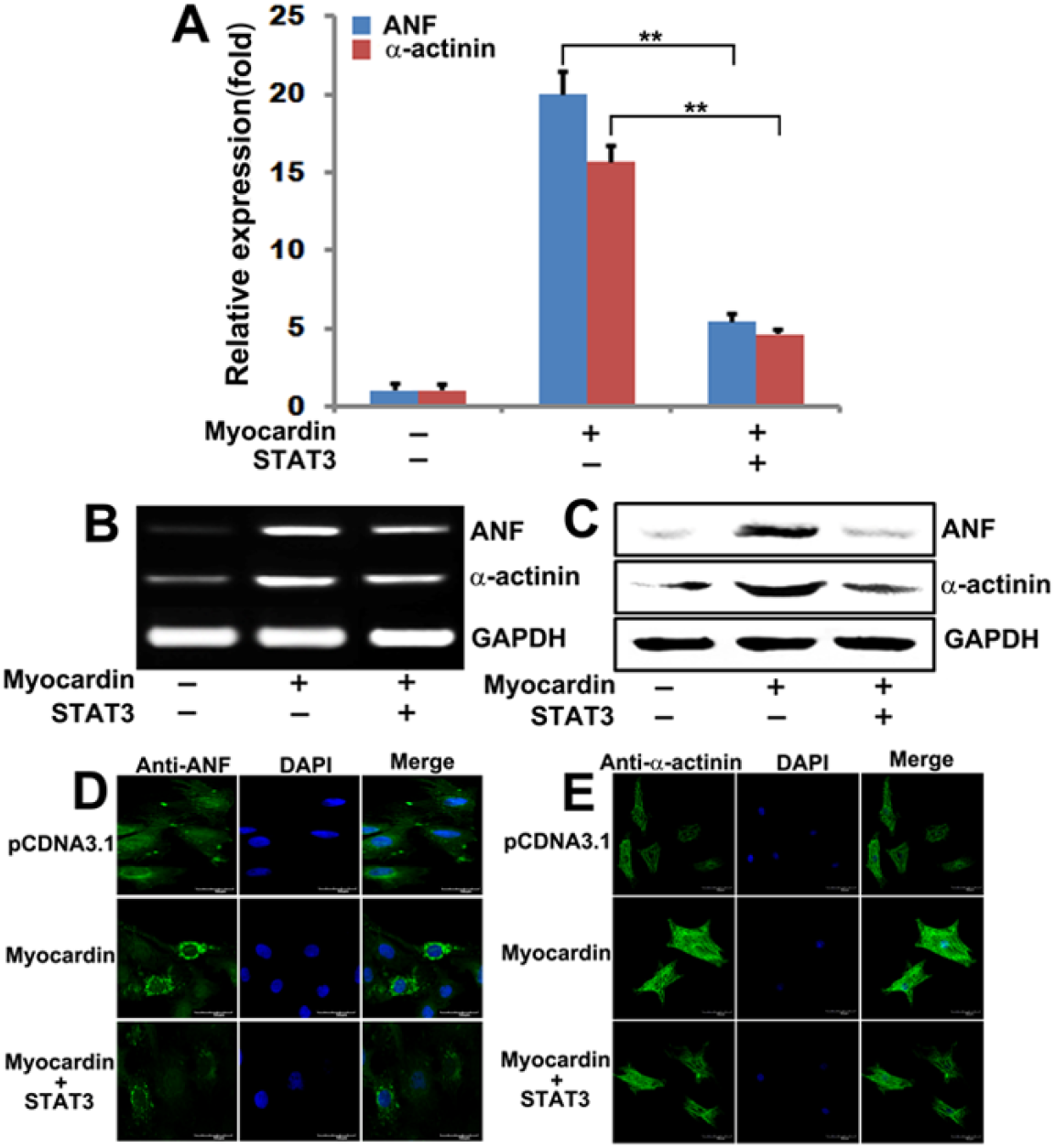
STAT3 suppresses Myocardin activation of cardiac genes. **A and B** qPCR and RT-PCR analysis of transfected with Myocardin or Myocardin/STAT3 for 48 h in cardiomyocytes. GAPDH served as loading control. (**, *p* < 0.01) Data are expressed as the mean□(±□SEM) with N□=□3 biological replicates in each group. All data are analysed using paired t-test. **C** Western blot analysis of transfected with Myocardin or Myocardin/STAT3 for 48 h in cardiomyocytes. GAPDH served as loading control. **D** The representative image shows ANF and α-actinin expression in cardiomyocytes transfected with Myocardin or Myocardin/STAT3 for 48 h. The left panels (green) show anti-ANF or α-actinin antibody reactivity to demonstrate gross morphology. The middle panels (blue) show the DAPI staining for nuclei. The right panels show double immunostained for ANF or α-actinin and nuclei. Scale = 50μm.

**Fig. 2.**
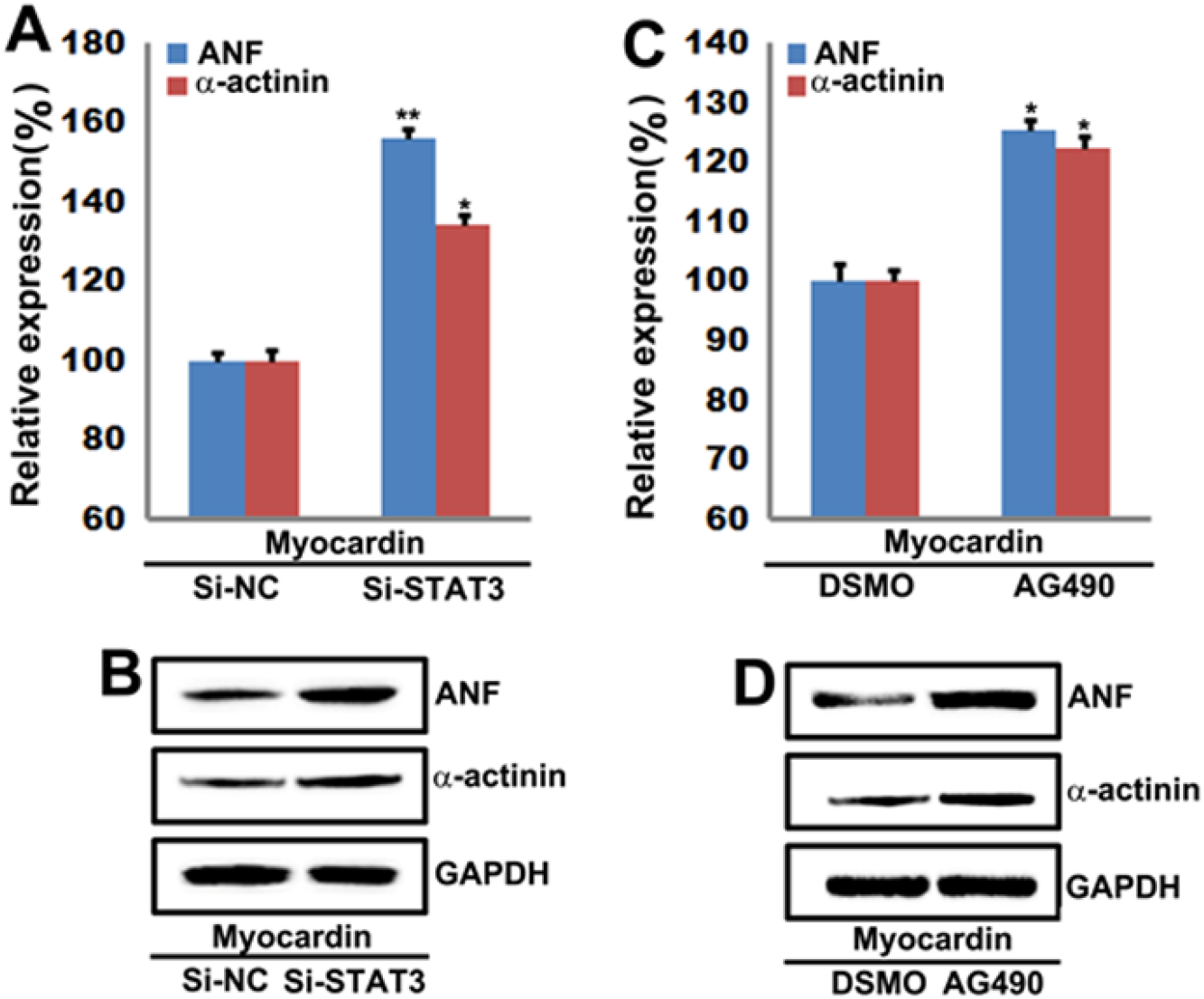
Silenced STAT3 and inhibited phosphorylation of STAT3 via AG490 can relieve STAT3 inhibiting Myocardin activation of cardiac genes. **A** qPCR analysis to detect the mRNA lever of ANF and α-actinin in cardiomyocytes transfected with Myocardin and then treated with siSTAT3 for 48 h. GAPDH served as loading control. (**, *p*< 0.01, *, *p*< 0.05). **B** Western blot analysis to detect the protein lever of ANF and α-actinin transfected with Myocardin and then treated with siSTAT3 for 48 h in cardiomyocytes. GAPDH served as loading control. **C** qPCR analysis to detect the mRNA lever of ANF and α-actinin in cardiomyocytes transfected with Myocardin and then treated with AG490 for 24 h. GAPDH served as loading control. (**, *p* < 0.01, *, *p* < 0.05). **D** Western blot analysis to detect the protein lever of ANF and α-actinin transfected with Myocardin and then treated with AG490 for 24 h in cardiomyocytes. GAPDH served as loading control. Data are expressed as the mean□ (± SEM) with N□=□3 biological replicates in each group. All data are analysed using paired t-test.

The JAK inhibitor, AG490 was used to attenuated endothelial STAT3 activation. Our results showed that AG490 significantly relieved the mRNA and protein expression of STAT3 suppressed Myocardin activation of cardiac genes in the presence of Myocardin in cardiomyocytes (Fig. 2C, D).

Taken together, these results indicate that STAT3 antagonizes the activation of myocardial genes by Myocardin.

### STAT3 affects the expression of Myocardin

To investigate the mechanism of STAT3 suppressed Myocardin activation of cardiac genes, qPCR assays were performed after overexpression of STAT3 in primary neonatal cardiomyocytes. Our data showed that overexpressing STAT3 did not affect the mRNA level of Myocardin (Fig. 3A).

**Fig. 3.**
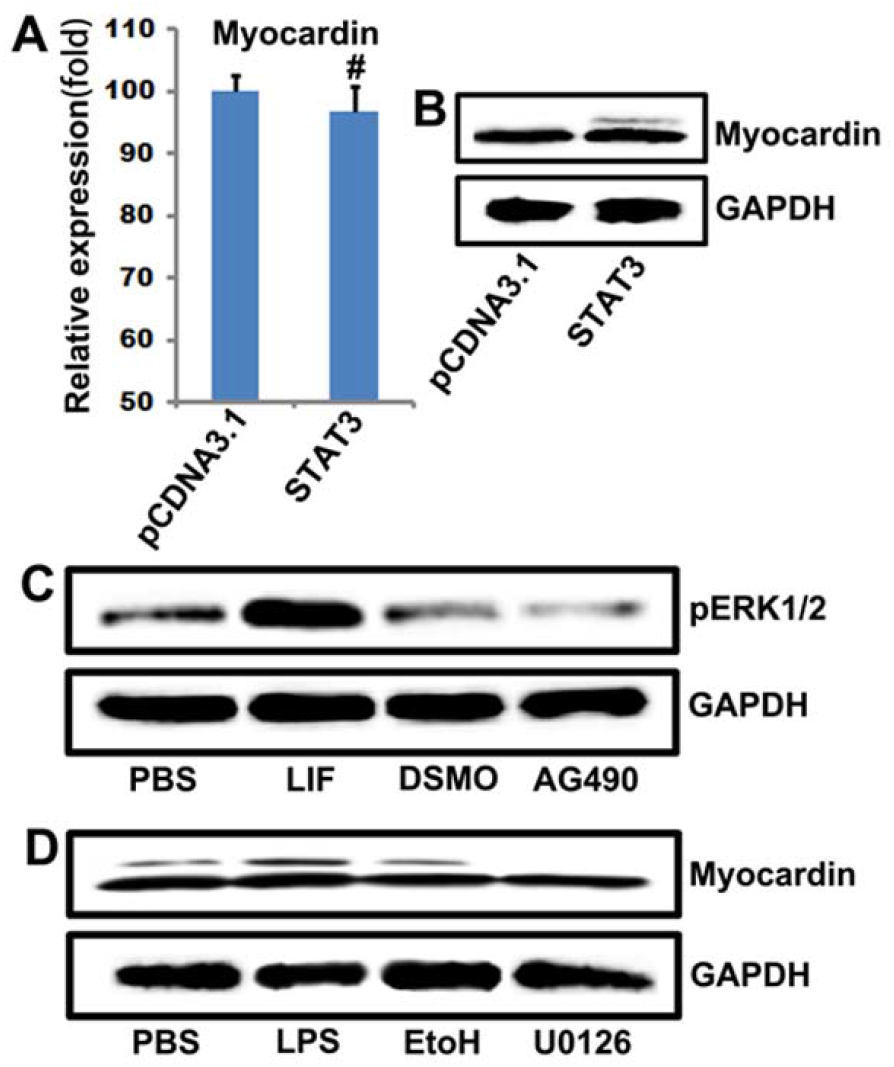
STAT3 affects the expression of Myocardin. **A** and **B** qPCR and Western blot analysis to detect expression of Myocardin in cardiomyocytes transfected with STAT3 for 48 h. GAPDH served as loading control. (^#^, *p*> 0.05) Data are expressed as the mean□ (± □ SEM) with N□=□3 biological replicates in each group. All data are analysed using paired t-test. **C** Western blot analysis of the phosphorylated ERK1/2 treated with LIF or AG490 for 24 h in cardiomyocytes. **D** Western blot analysis of the expression of Myocardin treated with LPS or U0126 for 24 h in cardiomyocytes.

Then we tested whether STAT3 could regulate the stability of Myocardin protein. The primary neonatal cardiomyocytes were transfected with a STAT3 plasmid for 48 h. The expression of endogenous Myocardin has a litter decreased by overexpressing STAT3 (Fig. 3B) and overexpressing STAT3 maybe phosphorylate the protein of Myocardin (see the top band in Fig. 3B). There have researches shown that phosphorylation of Myocardin by extracellular regulated protein kinases (ERK1/2) and phosphorylation of Myocardin by ERK1/2 reduces its induction of smooth muscle gene transcription.(17) Our results have shown that the STAT3 activator LIF promotes the phosphorylation of ERK but the STAT3 inhibitor AG490 inhibits ERK phosphorylation (Fig. 3C). Interestingly, the ERK activator lipopolysaccharide (LPS) promotes Myocaridn phosphorylation but the ERK inhibitor U0126 can inhibit the phosphorylation of Myocaridn (Fig. 3D). Thus, STAT3-ERK1/2-Myocardin forms a negative feedback loop regulating cardiomyocyte hypertrophy.

### STAT3 interacts directly with Myocardin

Next, we investigated whether myocardiin and STAT3 protein interact with co-immunoprecipitation assays with transfected COS7 cells. As shown in Fig. 7, Myocardin specifically interacted with STAT3. To locate the domains of cardiomycin and STAT3 that mediate interactions, a series of Myocardiin and STAT3 deletion mutant proteins were used by coimmunoprecipitation (Fig. 4A, C). Myocardin protein deletion mutants lacking N-terminal residues up to amino acid 194 interact with STAT3, while mutant proteins lacking first 268 amino acids fail to interact with STAT3. The ability to interact with STAT3 was also retained when the protein of the 265 C-terminal residue amino acid was deleted. Thus, STAT3 interacted with a distinct region near the N terminus (between amino acids 194 and 265, this is basic domain) (Fig. 4B). Myocardiin was confirmed to interact with SRF through a basic and glutamine-rich domain near the N-terminus.(18, 19) Because the STAT3-binding region of Myocardin overlapped the SRF-binding region, we considered the possibility that STAT3 may inhibit Myocardiin activity by replacing it with SRF.. The ability of STAT3 to inhibit the activity of Myocardin depended ANF-luciferase reporter activity also reduces the potential replacement mechanism of STAT3 inhibition. (Fig. 6).

**Fig. 4.**
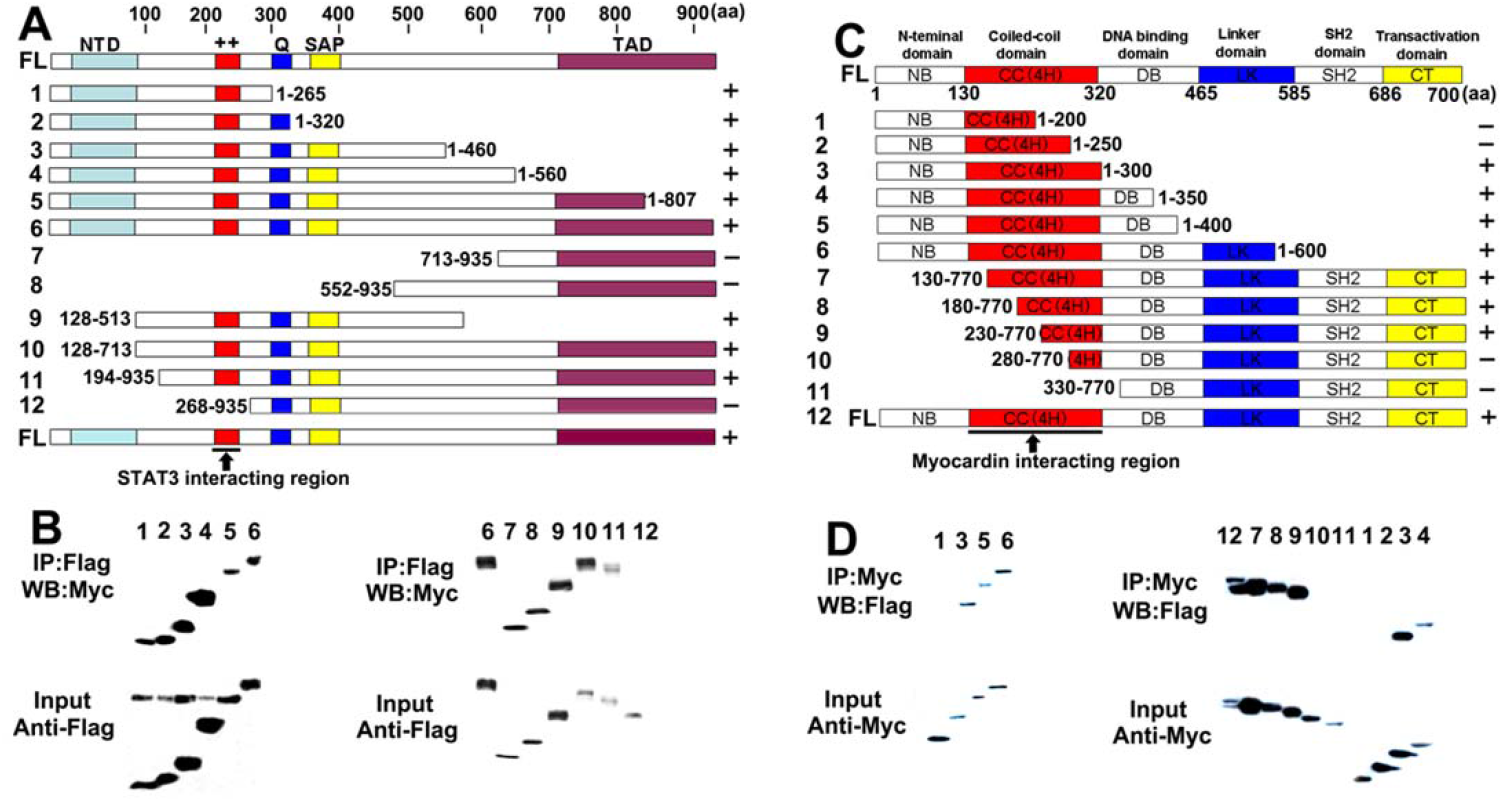
Direct interaction of Myocardin and STAT3 and mapping of the domains that mediate their interaction. **A** Schematic diagram of Myocardin and the mutant forms used to map the STAT3-binding domain. **B** Coimmunoprecipitation assays. COS7 cells were transiently transfected with expression vectors encoding Flag-tagged Myocardin deletion mutant proteins and Myc-tagged STAT3. Flag-tagged Myocardin deletion mutant proteins were immunoprecipitated (IP) from cell lysates with a monoclonal anti-Flag antibody, and coimmunoprecipitating STAT3 was detected by immunoblotting (IB) with a monoclonal anti-Myc antibody (top parts). The membrane was reprobed with anti-Flag antibody to reveal the total amount of Flag-tagged Myocardin proteins (bottom parts). **C** Schematic diagram of STAT3 and the mutant forms used to map the Myocardin-binding domain. **D** Coimmunoprecipitation assays. COS7 cells were transiently transfected with expression vectors encoding Myc-tagged STAT3 deletion mutant proteins and Flag-tagged Myocardin. Myc-tagged STAT3 deletion mutant proteins were immunoprecipitated (IP) from cell lysates with a monoclonal anti-Flag antibody, and coimmunoprecipitating Myocardin was detected by immunoblotting (IB) with a monoclonal anti-Flag antibody (top parts). The membrane was reprobed with anti-Myc antibody to reveal the total amount of Myc-tagged STAT3 proteins (bottom parts).

With a series of STAT3 deletion mutant proteins, we found that a mutant protein lacking the 300 sequence of amino acid N still retains the ability to interact with Myocardin (Fig. 4D). However, this interaction was completely eliminated when the amino acid 250 was further deleted, suggesting that the region between residues 250 and 300 is essential for STAT3 to interact with Myocardin. C-terminal deletion results indicate that residues downstream of amino acid 280 are not required to interact with Myocardiin (Fig. 4D). We concluded that a Myocardin-binding domain lies between amino acids 250 and 280 (coiled-coil domain) of STAT3.

When a STAT3 deletion mutant protein is present, there is a direct correlation between its ability to interact with and inhibit its activity. Thus, STAT3 can interact with and inhibit Myocardin, and only those STAT3 deletion mutant proteins that contain amino acids 250 and 280. It was worth noting that the C-terminal region of STAT3 (containing a transactivation domain) was not necessary for inhibiting myocardiin activity (Fig. 4D). These results demonstrated a direct interaction between the amino acids 250 and 280 (coiled-coil domain) of STAT3 and amino acids 194 and 265 (basic domain) Myocardin.

### Mutation the coiled-coil domain of STAT3 and the basic domain of Myocardin affect the interaction of Myocardin and STAT3 and its transcriptional activation

This deletion mutant Myocardin basic domain [Myocardin (1-807)] abolished interactions between Myocardin and STAT3 but this mutant Myocardin Q-rich domain [Myocardin (1-807)] has interactions between STAT3. A construct containing this basic domain [Myocardin (1-807)] also bound STAT3 (Fig. 5A, B). These studies further support that the basic domain of Myocardin interacts in vitro with STAT3. Since a direct interaction between the amino acids 250 and 280 (coiled-coil domain) of STAT3 and amino acids 194 and 265 (basic domain) Myocardin, we have mutated site-directed the amino acids (KK-NQ) of Myocardin basic domain (Mutation-Myocardin) (Fig. 5C) or the amino acids (DDE-GSQ) of STAT3 coiled-coil domain (Mutation-STAT3) (Fig. 5D).

**Fig. 5.**
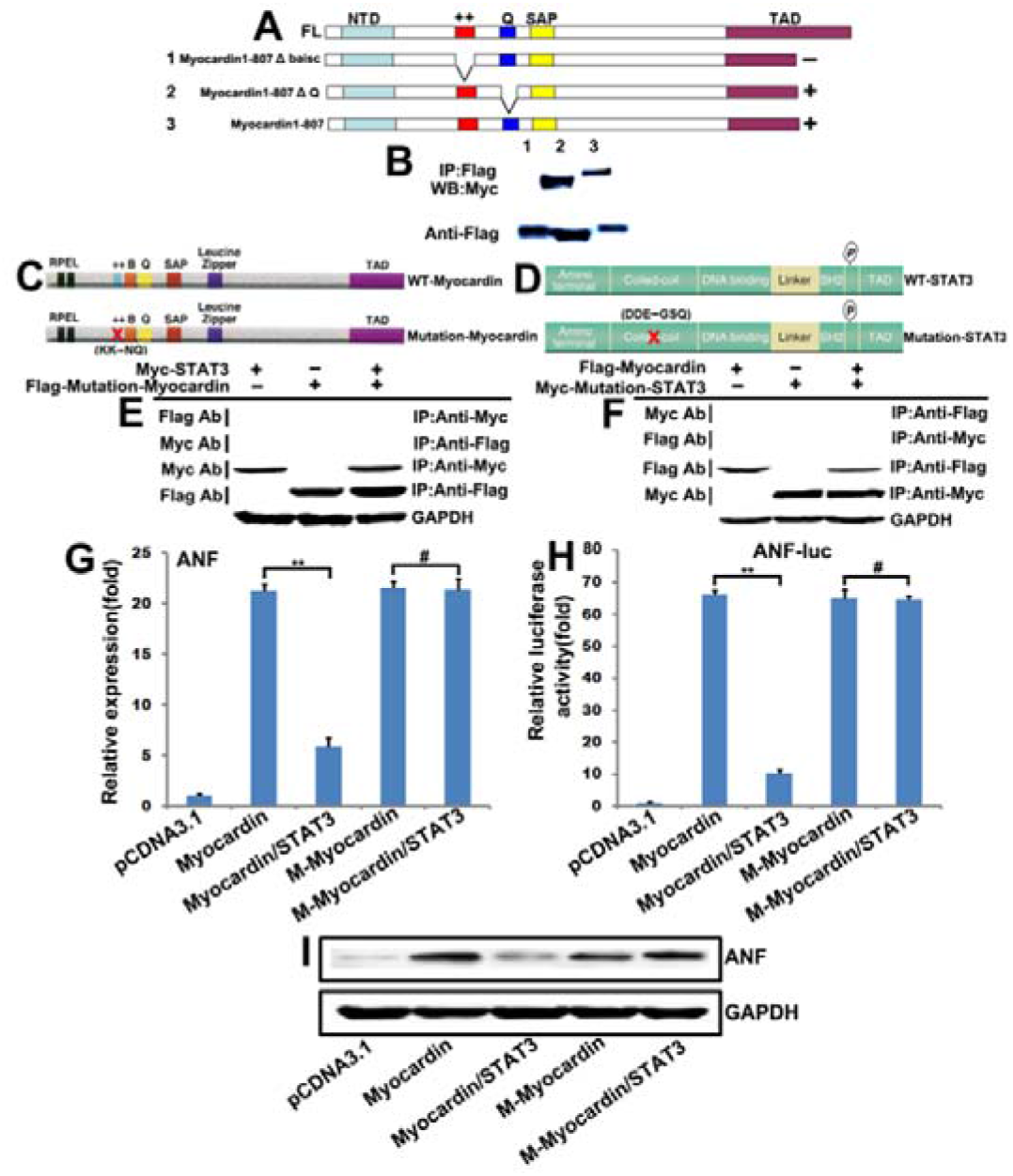
Mutation the coiled-coil domain of STAT3 and the basic domain of Myocardin affect the interaction of Myocardin and STAT3 and its transcriptional activation. **A** Coimmunoprecipitation assays. COS7 cells were transiently transfected with expression vectors encoding Flag-tagged Myocardin deletion mutant (basic domain and SAP domain) proteins and Myc-tagged STAT3. Flag-tagged Myocardin deletion mutant proteins were immunoprecipitated (IP) from cell lysates with a monoclonal anti-Flag antibody, and coimmunoprecipitating STAT3 was detected by immunoblotting (IB) with a monoclonal anti-Myc antibody (top parts). The membrane was reprobed with anti-Flag antibody to reveal the total amount of Flag-tagged Myocardin proteins (bottom parts). **B** and **C** Schematic of mutated site-directed the amino acids (KK-NQ) of Myocardin basic domain (Mutation-Myocardin) and mutated site-directed the amino acids (DDE-GSQ) of STAT3 coiled-coil domain (Mutation-STAT3). **E** and **F** Coimmunoprecipitation assays in mutated site-directed the amino acids (KK-NQ) of Myocardin basic domain (Mutation-Myocardin) and STAT3 or mutated site-directed the amino acids (DDE-GSQ) of STAT3 coiled-coil domain (Mutation-STAT3) and Myocardin. **G, H** and **I** QPCR, Luciferase assay and Western blot analysis to detect expression of ANF in cardiomyocytes transfected with Mutation the coiled-coil domain of STAT3 and Myocardin or the basic domain of Myocardin and STAT3 for 48 h. GAPDH served as loading control (**, *p* < 0.01, ^#^, *p* > 0.05) Data are expressed as the mean□(±□SEM) with N□=□3 biological replicates in each group. All data are analysed using paired t-test.

Coimmunoprecipitation assays to test the interaction mutated site-directed the amino acids (KK-NQ) of Myocardin basic domain (Mutation-Myocardin) and STAT3 or mutated site-directed the amino acids (DDE-GSQ) of STAT3 coiled-coil domain (Mutation-STAT3) and Myocardin. Our data show that Mutation-Myocardin has no interaction with STAT3 (Fig. 5E) and Mutation-STAT3 has no interaction with Myocardin (Fig. 5F). Taken together, these data indicate that the interaction between Myocardin and STAT3 is produced by the positive charge of the amino acids KK of the Myocardin basic domain and the negative charge of the amino acids DDE of the STAT3 coiled-coil domain.

To investigate the role of transcription activation and expression of ANF after mutation the coiled-coil domain of STAT3 and the basic domain of Myocardin, the qPCR and luciferase reporter and western bolt assay were used to test the transcription activation and expression of ANF. As shown in Fig. 5G, H and I, overexpressed Myocardin will strongly active the expression of cardiac gene (ANF) but the cotransfected Myocardin and STAT3 decreased Myocardin-induced cardiac gene expression compared with that in transfected Myocardin cells, Mutation-STAT3 significantly relieved the mRNA and protein expression of STAT3 suppressed Myocardin activation of ANF in the presence of Myocardin in cardiomyocytes. More importantly, Mutation-Myocardin significantly relieved the mRNA and protein expression of STAT3 suppressed Myocardin activation of ANF in the presence of STAT3 in cardiomyocytes. Thus, STAT3 inhibits Myocardin induced cardiac hypertrophy gene expression through their interaction, mutation the interaction of the site, this effect has disappeared.

### STAT3 interferes with the SRF–Myocardin interaction

In order to understand the molecular mechanism by which STAT3 inhibits myocardiin function, we asked whether STAT3 can interfere with myocardiin-SRF interactions. To investigate this, we transfected cells with Myc-Myocardin and HA-SRF and followed their binding after silencing (Figure 6A) or over-expressing STAT3 (Fig. 6B). The former condition was significantly enhanced, while the latter significantly reduced the binding of SRF to Myocardin. Indeed, Δcoiled-coil retained substantial ANF promoter-inducing activity (Fig. 6D), whereas it combination with STAT3 was greatly reduced (Fig. 6C). Importantly, Δcoiled-coil was much less sensitive to the inhibitory action of STAT3 than the WT (Fig. 6D). These findings suggest that the key mechanism for STAT3-mediated ANF promoter inhibition is the binding of STAT3 to Myocardin.

**Fig. 6.**
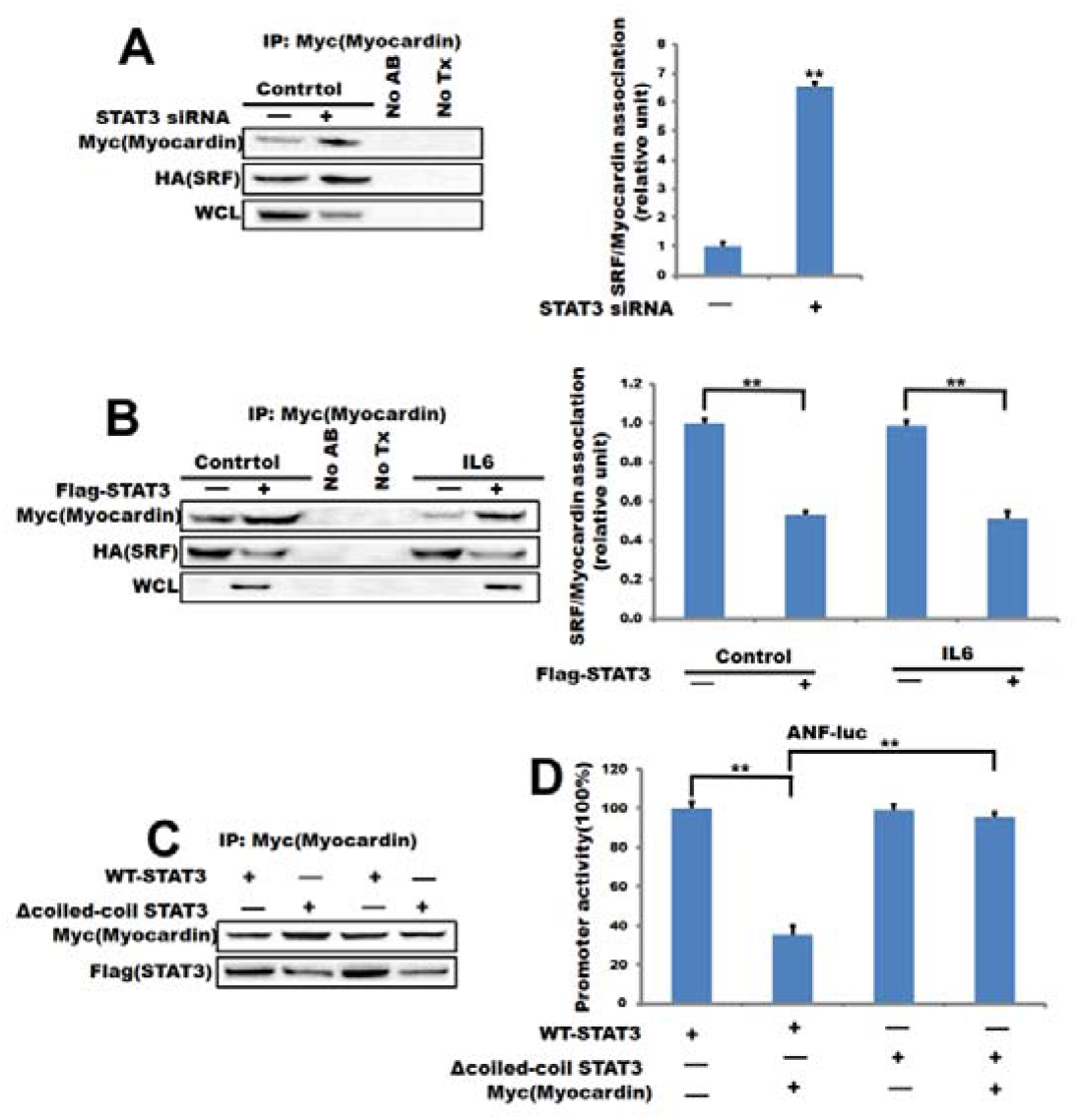
STAT3 interferes with Myocardin–SRF interaction. **A** Cells were transfected with Myc-Myocardin and HA-SRF along with NC or STAT3 siRNA. Association of Myocardin and SRF was analyzed by coimmunoprecipitation. STAT3 silencing was detected from whole cell lysates (WCL). Controls for the immunoprecipitation were reaction without antibody (No AB) or Myc transfection (No Tx). (right) Densitometric analysis of three experiments is shown. **B** Myc-Myocardin and HA-SRF were cotransfected with empty vector or STAT3. Myocardin was immunoprecipitated with anti-Myc antibody as in A from control or LCM-treated (1 h) cells. **C** Coimmunoprecipitation shows decreased association of STAT3 to Δcoiled-coil compared with WT. **D** Coiled-coil domain mutant shows reduced sensitivity to inhibition by STAT3. Cells were transfected with ANF-Luc, Δcoiled-coil, or WT-Myocardin along with empty vector or STAT3. Luciferase assay was performed 48 h later. Results are normalized to the control (top; fold increase over control) or expressed as a percentage of the maximal effect of the given Myocardin construct (bottom). **, *p* < 0.01. Error bars indicate mean ± SEM.

### Myocardin and STAT3 promote the transactivity of ANF depending on CArG box

Myocardin coordinate and regulate the expression of a variety of CArG-dependent cardiac genes. Previously, it was found that incorrect expression of cardiac proteins in Xenopus cap analysis was not sufficient to induce beating or cardiac-like coordinated contractions.(20) Although Myocardiin does not directly bind to DNA, the binding of SRF to the CArG box (CC (A/T)_6_GG) will cause a sharp bend in the DNA, and this bend may change with the change in base composition in the entire CArG box. (21) All STATs bind to similar SIE/GAS DNA sequences that contain the TT and AA base tandems, separated in STAT3 by 4 to 6 bases (5’-TTN_4-6_AA-3’).(22, 23) By bioinformatics analysis, we found that the ANF promoter has two CArG box sites and a GAS-like site (Fig. 7A). To further determine whether STAT3 affects the transactivation of Myocardial-mediated ANF promoters and to investigate whether this process depends on the CArG box, The ANF gene promoter luciferase reporter plasmid was constructed to contain a deleted or mutated CArG box. (Fig. 7A, B). Our data indicate that Myocardial-mediated ANF promoter activity is abolished in vitro when the CArG box is deleted or mutated in the ANF promoter, while nearby CArG boxes play an important role in Myocardial-mediated ANF promoter activity (Fig. 7C, D). The above experimental data show that the transactivation of ANF by Myocardin depends on the CArG box.

**Fig. 7.**
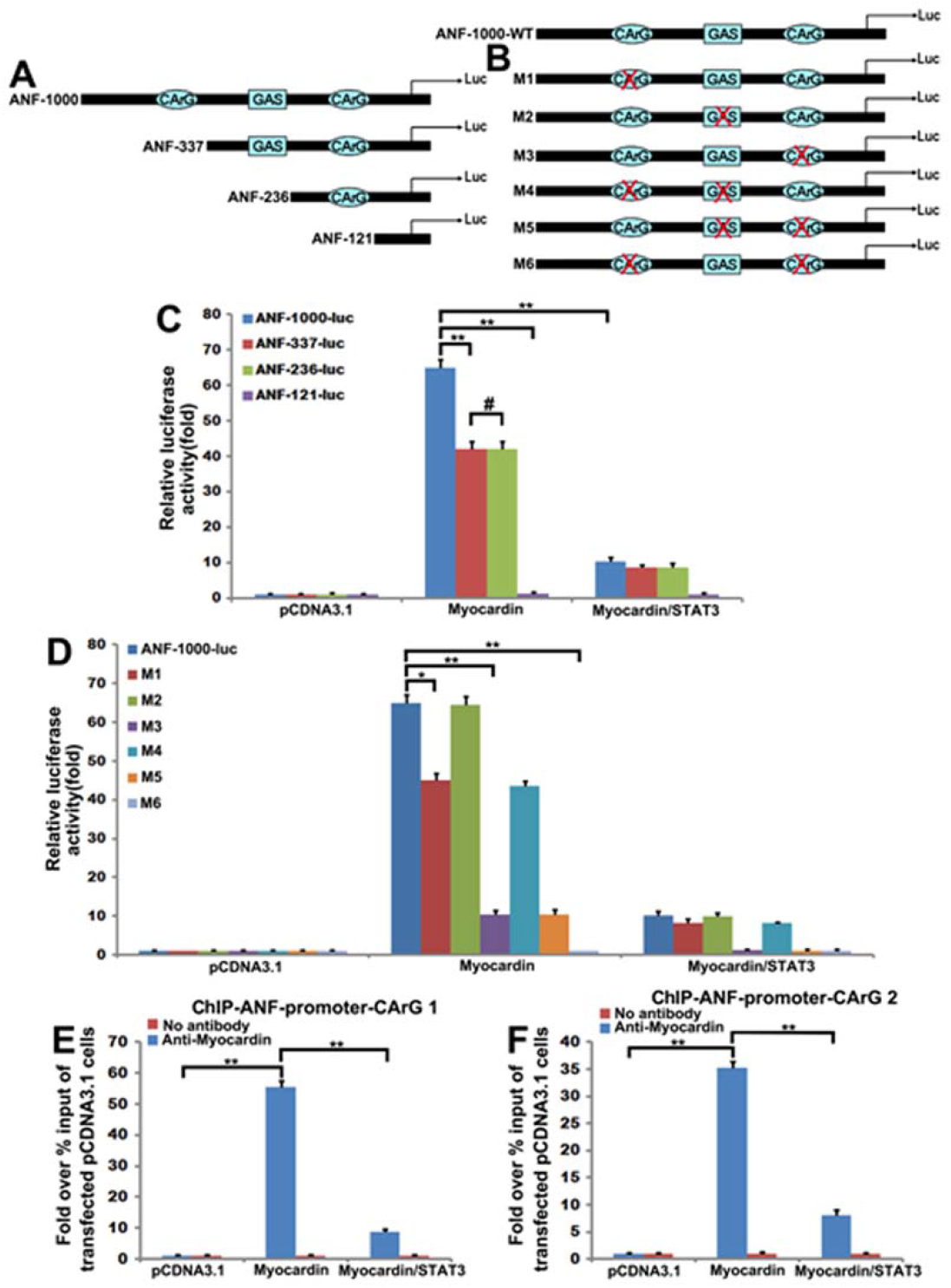
STAT3 inhibits Myocardin-induced the transactivity of ANF depending on CArG box. **A** Schematic of the −1000 ANF promoter, containing CArG box element was linked to a luciferase reporter. Mutation or truncations that remove the CArG box element. −337, −236 and −121 ANF promoter, a truncated promoter. **B** The −1000 ANF promoter with mutations in CArG box. **C** Cardiomyocytes were transfected with the Wild-Type-1000 ANF promoter, or a truncated promoter −337, −236 or −121 and transfected with Myocardin or STAT3 for 48 h. Then the luciferase reporter assays were used to test the transactivity of ANF. **D** Cardiomyocytes were transfected with the Wild-Type-1000 ANF promoter, or −1000 ANF promoter with mutations in CArG box and transfected with Myocardin or STAT3 for 48 h. Then the luciferase reporter assays were used to test the transactivity of ANF. (**, *p* < 0.01, *, *p* < 0.05). **E and F** Cardiomyocytes were transiently transfected with a Myocardin, Myocardin /STAT3 or a control vector (pCDNA3.1) 48 h, and ChIP assays were performed by PCR with primers associated with the genes for ANF as described in Materials and Methods. Sheared DNA/protein complexes were immunoprecipitated by using an anti-Flag-Myocardin Ab. Then, PCR was carried out to detect the endogenous CArG regions in immunoprecipitated chromatin fragments. The amount of DNA in each sample (2% input) is shown at the second land. Immunoprecipitations were performed without primary antibody (No Ab) as a control and IgG as a negative control (**, *p*< 0.01). Data are expressed as the mean□(±□ SEM) with N□=□3 biological replicates in each group. All data are analysed using paired t-test.

As shown in Fig. 7C and 7D, overexpressing Myocardin strongly activated the expression of ANF-1000-luc promoter. However, coexpression Myocardin and STAT3 repressed Myocardin-dependent ANF transcriptional activity compared with that in transfected Myocardin cells. These data demonstrate that STAT3 serves as a negative regulator of cardiac muscle differentiation in the presence of Myocardin through downregulation of Myocardin activity.

To further confirm the specific binding of Myocardin to ANF promoter, ChIP assays were performed in cardiomyocytes transfected with Myocardin, Myocardin /STAT3 or vector. Immunoprecipitation of cross-linked chromatin with specific antibodies against Myocardin or without antibodies (as negative control). The precipitated chromatin DNA was then purified and PCR amplified using specific primers across the CArG box in the ANF promoter. As shown in Fig. 7E and 7F, the comparison (IgG) and the negative control (No Ab) did not show any PCR signals. Myocardiin can be combined with the CArG box of the ANF promoter (both near (CArG 1) and long-range (CArG 2) CArG boxes). Myocardiin binds to the near CArG box more than the ability to bind to the far CArG box in the ANF promoter (Fig. 7E, F). More importantly, STAT3 has the effect of inhibiting Myocardiin. (Fig. 7E, F). These data indicate that STAT3 is a potent nuclear factor that inhibits the transactivation of the Myocardium by disrupting the formation of the SRF/Myocardin/CArG complex in vivo and in vitro.

### Knockdown stat3 inhibits Myocardin knockdown-mediated cardiac hypertrophy in zebrafish

To further validate the conclusions in vitro and the effects of Myocardin and STAT3 interactions on cardiac development, Myocardin-MO, STAT3-MO and control-MO targeting ATG were synthesized and microinjection into the zebrafish embryos. The developmental process of the zebrafish heart is observed by the expression of the cardiac development marker gene CMCL2. The results show that the heart cannot be circulated normally when STAT3 is knocked down. While co-injecting Myocardin-MO and STAT3-MO, most of the fish’s heart showed near normal cyclization, but there were still some malformations (Fig. 8A). We also characterized cardiac hypertrophy by detecting the ANF gene. Microinjection of STAT3-MO alone induced cardiac hypertrophy. When we injected Myocardin-MO and STAT3-MO together, cardiac hypertrophy was alleviated (Fig. 8B). In addition, significant up-regulation of ANF expression could be clearly observed when knocking down STAT3 (Fig. 8B), which confirmed our conclusions in primary cardiomyocytes. These results indicated that STAT3 could antagonize Myocardin-mediated cardiac hypertrophy in vivo.

**Fig. 8.**
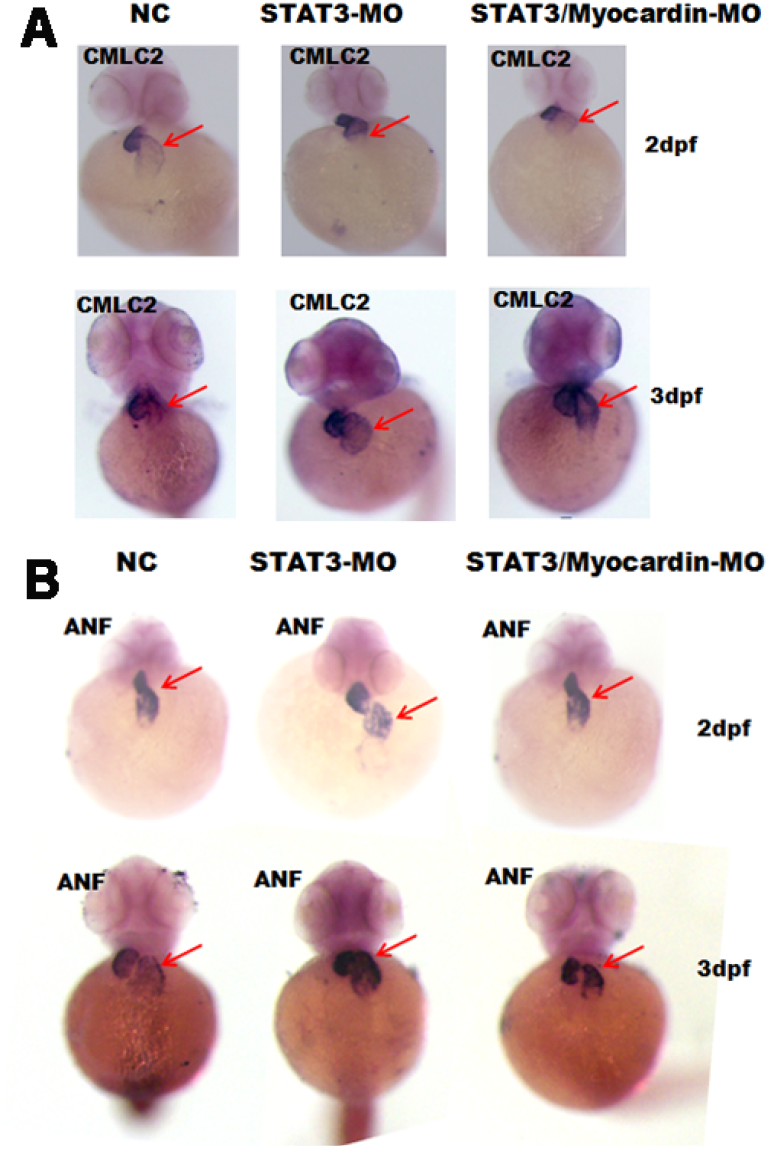
STAT3 antagonizes Myocardin-mediated cardiac hypertrophy. Microinjection of Myocardin-MO, STAT3/Myocardin-MO and control-MO separately into the zebrafish embryos, and then the zebrafish embryos were subjected to in situ hybridization of the entire embryo. **A** Expression of CMLC2 2 days (upper) and 3 days (bottom) after fertilization. (n>30). **B** Expression of ANF 2 days (upper) and 3 days (bottom) after fertilization. (n>30).

## Discussion

Myocardin, which plays a key role in inducing cardiac hypertrophy and fetal cardiac genetic programs, as a co-activator of SRF,(1). However, the molecular mechanism in regulating the stability and activity of Myocardin is unclear. Despite recent evidence that STAT3 plays a cardioprotective role in the heart, there is no information on the role of STAT3 in Myocardial-induced myocardial hypertrophy. This study revealed the relationship between Myocardiin and STAT3 in the development of cardiac hypertrophy.

Our study has shown that Myocardin and STAT3 have physically interaction by co-immunoprecipitation. More importantly, our data demonstrated a direct interaction between the amino acids 250 and 280 (coiled-coil domain) of STAT3 and amino acids 194 and 265 (basic domain) Myocardin (Fig. 4). So that we hypothesized that STAT3 was a coactivator for Myocardin, the regulator of cardiomyocyte hypertrophy. Interestingly, we find that co-expession of Myocardin and STAT3 can decrease Myocardin-induced cardiac gene expression compared with that in transfected Myocardin alone in cardiomyocytes (Fig. 1). Silenced STAT3 via siSTAT3 can relieve STAT3 inhibiting Myocardin activation of cardiac genes (Fig. 2A, B). Microinjection of STAT3-MO alone into zebrafish induced cardiac hypertrophy. When we injected Myocardin-MO and STAT3-MO together, cardiac hypertrophy was alleviated (Fig. 8B). In addition, significant up-regulation of ANF expression could be clearly observed when knocking down STAT3, which confirmed our conclusions in primary cardiomyocytes. The influence that AG490 could block the phosphorylation of JAK2 would restrain the phosphorylation of STAT3. AG490 would weak the effect of STAT3 inhibit Myocardin by restraining the phosphorylation of STAT3 (Fig. 2C, D). Mutation the coiled-coil domain of STAT3 and the basic domain of Myocardin affect the interaction of Myocardin and STAT3 and Myocardin activation of cardiac genes transcriptional activation (Fig. 5G, H and I).

We also provide evidence that far-CArG (CArG2) and near-CArG (CArG1) in the ANF promoter are essential for promoter function in cardiomyocytes. But the function of near-CArG is more powerful than the far-CArG (Fig. 7C, D). These results indicate that this spacing of the paired CArG elements contributes to the synergistic combination of SRF and its coactivator Myocardin with the ANF promoter. STAT3 inhibits Myocardial activation of CArG-dependent cardiac gene expression. The effect mediated by physical interruption of the Myocardin/SRF/CArG ternary complex (Fig. 7E, F). Our data demonstrate that STAT3 can behave as a negative transcriptional cofactor in a Myocardin-dependent manner to inhibit cardiomyocyte hypertrophy.

Why does STAT3 inhibit Myocardin transactivity? Firstly, STAT3 can directly inhibit Myocardin-dependent transcriptional activity, qPCR assays were performed in transfected STAT3 primary neonatal cardiomyocytes. But our data show that overexpressing STAT3 did not affect the mRNA level Myocardin. The expression of endogenous Myocardin has a litter decreased by overexpressing STAT3 (Fig. 3B) and overexpressing STAT3 maybe phosphorylate the protein of Myocardin (see the top band in Fig. 3B). There have researches shown that phosphorylation of Myocardin by ERK1/2 and phosphorylation of Myocardin by ERK1/2 reduces its induction of smooth muscle gene transcription.(17, 24) Our results have shown that the STAT3 activator LIF promotes the phosphorylation of ERK but the STAT3 inhibitor AG490 inhibits ERK phosphorylation (Fig. 3C). Interestingly, the ERK activator LPS promotes Myocaridn phosphorylation but the ERK inhibitor U0126 can inhibit the phosphorylation of Myocaridn (Fig. 3 D). The molecular mechanism may be the phosphorylation modification of Myocardin by STAT3. And the phosphorylation modification of Myocardin might be prone to decompound by ubiquitin. Recently, Protein inhibitor of activated STAT 1(PIAS1) has been reported to regulate Myocardin transactivity via its ligase induced sumoylation. These results indicate that PIAS1 and small ubiquitin-like modifier 1 (SUMO1) enhance Myocardiin activity through SUMO-modified Myocardial proteins..(25, 26) Protein inhibitor of activated STAT represents a family of proteins originally identified through interaction with cytokine-induced STAT. (27) PIAS1 binds to STAT1 and inhibits the transcriptional activity of STAT1 by blocking its DNA-binding activity.(28, 29) STAT3 and STAT1 form complexes in response to LIF stimulation.(30) Thus, it is also possible that STAT3 inhibits Myocardin transactivity by inhibiting SUMO modification of Myocardin by PIAS1 and SUMO1. The specific binding site that both SUMO modification of Myocardin by PIAS1/SUMO1 and ubiquitin modification is the site KXE (IKQE). The competition between SUMO modification and ubiquitin modification of Myocardin at the same site KXE (IKQE) regulates Myocardin transactivity.

Taken together, these data demonstrate that STAT3 serves as a negative regulator of Myocardin activity to regulate cardiomyocyte hypertrophy.

## Conclusion

The N-terminus of STAT3 directly binds to the basic domain of myocardin and inhibits the transcriptional activity of myocardin-mediated cardiac-specific genes ANF and α-actinin, thereby inhibiting their expression, and further inhibit myocardin-mediated cardiac hypertrophy in vivo. The STAT3-Myocardin interaction identifies a site of convergence for nuclear hormonereceptor-mediated and cardiac-specific gene regulation and suggests a possible mechanism for the cardiac protective effects.

## Abbreviations

STAT3: signal transducer and activator of transcription 3;
SRF: serum response factor;
IL-6: interleukin-6;
LIF: leukemia inhibitory factor;
DMEM: Dulbecco’s modified Eagle’s medium;
DAPI: 4’,6’-diamidino-2-phenylindole;
RT-PCR: semi-quantitative reverse-transcription polymerase chain reaction;
qPCR: real time quantitative polymerase chain reaction;
GAPDH: Glyceraldehyde-3-phosphate dehydrogenase;
PAGE: sodium dodecyl sulfate-polyacrylamide gel electrophoresis;
ChIP: Chromatin Immunoprecipitation;
JAK: Janus kinase;
LPS: lipopolysaccharide;
ERK1/2: extracellular regulated protein kinases;
PIAS1: protein inhibitor of activated STAT 1;
SUMO1: small uniquitin-like modifier 1;
MO: Morpholino

## Ethics declarations

### Ethics approval and consent to participate

The research is approved by an ethical committee of Wuhan University of Science and Technology.

### Consent for publication

Not applicable.

### Availability of data and materials

All data, models, and code generated or used during the study appear in the submitted article. After all experiments were completed, they were executed using MS-222.

### Authors’ contributions

Xing-Hua Liao and Tong-Cun Zhang designed research; Jia-Peng Li, Yuan Xiang, Hui Li, You Huang and Chao Shen performed research; Jia-Peng Li, Yuan Xiang, Hui Li and Xing-Hua Liao analyzed data; Hui-Min Zhang, Xing-Hua Liao and Tong-Cun Zhang wrote the paper.

## Acknowledgements

Not applicable.

